# The biogeography of the stripped Venus clam, *Chamelea gallina*, in the Mediterranean Sea indicates limited gene flow and shows evidence of local adaptation

**DOI:** 10.64898/2026.02.08.704711

**Authors:** Benestan Laura, Marc Baeta, Carlos Saavedra, Marina Delgado, Silvia Falco, Miguel Rodilla, Luis Silva, Miriam Hampel, Ciro Rico

## Abstract

**Aim:** To assess how biogeographic barriers and environmental heterogeneity shape connectivity and local adaptation in the striped Venus clam (*Chamelea gallina*), a commercially exploited bivalve in the Mediterranean Sea.

**Location:** Northeast Atlantic (Gulf of Cádiz) and Mediterranean Sea (Alboran, Balearic, Tyrrhenian and Adriatic regions).

**Taxon:** *Chamelea gallina* (Bivalvia: Veneridae).

**Methods:** We analysed genome-wide single nucleotide polymorphisms (SNPs) from 226 individuals sampled across six regions (Gulf of Cadiz, Alboran Sea, Balearic Sea, Ebro Delta, Tyrrhenian Sea and Adriatic Sea) using a seascape genomic framework. Population structure was inferred using both putatively neutral and adaptive loci. Genotype–environment associations were tested against key oceanographic variables, including sea surface temperature, salinity and nutrient availability.

**Results:** Neutral loci revealed weak genetic differentiation, consistent with substantial gene flow across most of the species’ range. In contrast, putatively adaptive loci uncovered pronounced genetic structure that corresponded closely to major Mediterranean biogeographic regions, particularly the Adriatic Sea, the Gulf of Cadiz and western–central Mediterranean basins. Significant associations were detected between genetic variation and environmental gardients, with several candidate adaptive SNPs located within coding regions, suggesting functional responses to spatially heterogeneous conditions.

**Main conclusions:** Our results demonstrate that local adaptation can generate biologically meaningful population structure in *C. gallina* despite high levels of connectivity inferred from neutral markers. This decoupling between neutral and adaptive variation highlights the importance of integrating adaptive genomic information into biogeographic inference. Recognizing environmentally driven genetic differentiation is essential for defining robust management units and for improving the long-term sustainability and resilience of *C. gallina* fisheries under increasing anthropogenic pressure and climate change.

## 1 Introduction

Effective management of marine biota requires understanding spatial connectivity patterns and the environmental factors shaping dispersal (Benestan, 2020). Advances in molecular techniques now reveal genetic markers associated with local adaptation and connectivity (Leigh, Hendry, Vázquez-Domínguez, & Friesen, 2019). Integrating this knowledge into fisheries management enables strategies tailored to the ecological characteristics of each stock, supporting long-term sustainability (Snead & Clark, 2022). In this context, delineating biogeographic regions defined by distinct physicochemical, oceanographic and geomorphological features is key to guiding fisheries and conservation decisions (Lourie & Vincent, 2004).

The Mediterranean Sea is composed of five main regions: the Western, Central and Levantine basins, along with the Adriatic and Aegean Seas. Each region is characterized by a distinct oceanographic regime – for instance, the broad continental shelf of the Western Mediterranean, the shallow waters of the Adriatic Sea, the high temperatures of the Central Basin, the great depths of the Levantine Basin, and the relatively low salinity and temperature of the Aegean Sea (Arvanitidis et al., 2002). These regions corresponds to geomorphological sub-basins separated by straits and sills, whose unique physicochemical and oceanographic features often delineate biogeographical areas (Arvanitidis et al., 2002). Still, boundaries between biogeographic regions are not always clear, as environmental discontinuities affect taxa differently depending on dispersal ability and other life history traits (e.g., Galarza et al., 2009; Pascual, Rives, Schunter, & Macpherson, 2017). Within the Western Mediterranean and the Adriatic regions, additional biogeographic zones are defined by oceanographic fronts, environmental breaks and geomorphological barriers. Oceanographic processes such as prevailing currents, gyres and hydrodynamic discontinuities strongly influence population, acting as natural barriers that shape population structure (Boulanger et al., 2022; Galarza et al., 2009; Pascual et al., 2017; Sefc et al., 2020).

Oceanic fronts are key features in marine environments, acting as barriers to faunal exchange by creating sharp physical and biochemical variables discontinuities. The western Mediterranean, a well-known biodiversity hotspot with high endemism (Macpherson, 2002), is notably influenced by ocean circulation, especially the influx of Atlantic water through the Strait of Gibraltar (Millot, 2005). This interaction generates two major fronts: the Almeria-Oran Front (AOF) and the Balearic Front (BF). The AOF, situated approximately 400 km east of Gibraltar, is a semi-permanent thermohaline front extending to 300 meters depth, with density gradients comparable to those of the Gulf Stream (Tintoré, La Violette, Blade, & Cruzado, 1988). It has been shown to limit gene flow in several marine species (Galarza et al., 2009; Pascual et al., 2017). Furthermore, the BF is characterised by density differences of 0.5 σT units and modulates the transport of Atlantic waters towards the Balearic Islands. Together with the Balearic Current – an extension of the Northern Current progressing northeast over the northwestern slope of the Balearic Islands the BF delineates the Balearic sub-basin (Balbín et al., 2012). Like other semi-enclosed seas (Millot, 2005; Pinot, Tintore, & Gomis, 1994; Tintoré et al., 1988), the Balearic Sea offers a natural setting to examine how oceanographic fronts constrain connectivity (Galarza et al., 2009), in contrast to populations not subjected to such barriers (Pascual et al., 2017).

The Strait of Gibraltar, a narrow channel linking the Mediterranean Sea and the Atlantic Ocean, mediates unique water exchanges, while the AOF separates nutrient-rich from oligotrophic waters, shaping marine connectivity (Tintore, La Violette, Blade, & Cruzado, 1988). Atlantic waters near Gibraltar undergo recirculation driven by tidal currents and an underwater ridge, creating a cold-water zone that limits species migration between the Atlantic and Mediterranean (Sala et al., 2023). These oceanographic dynamics influence marine ecosystems by modifying species distribution and abundance (Marie et al., 2016; Nantón et al., 2017; T. Patarnello, F. A. Volckaert, & R. Castilho, 2007). Acting as a semi-permanent barrier to gene flow, the AOF has been associated with pronounced genetic differences among populations from the Atlantic, Alboran Sea, and north-western Mediterranean across a wide range of marine taxa (Pascual et al., 2017; Sánchez-Garrido & Nadal, 2022). The sharp environmental contrasts on either side of the AOF reinforce these genetic breaks.

The Adriatic Sea, covering 5.5% of the Mediterranean’s surface area, is a distinct biogeographic region (Sefc et al., 2020) with contrasting coastlines - a sandy western shore and a steep eastern one - with winter temperatures dropping below 10°C in the North. The sea is relatively shallow, with three-quarters of the seabed under 200 meters, and subdivided into North, Central, and South Adriatic sub-regions, each with specific depth, temperature, and biodiversity patterns (Bianchi et al., 2012). The North Adriatic, characterized by depths under 40 meters, is influenced by freshwater inputs, particularly from the Po River, and by two main recirculation gyres (Poulain, 2001; Ursella, Poulain, & Signell, 2006). This region sustains high productivity due to nutrient enrichment from freshwater discharges. In contrast, the Central and South Adriatic are generally less productive. Ocean circulation is further shaped by seasonal wind patterns, which influence larval dispersal and species distribution (Sefc et al., 2020). In comparison, the Tyrrhenian Sea is much deeper (> 2000 meters), with more stable salinity, lower productivity, and unique coralligenous habitats. Both regions are vulnerable to anthropogenic pressures, especially agricultural runoff and industrial activities, which degrade water quality (Cognetti, Lardicci, Abbiati, & Castelli, 2000).

Bivalves, including commercially valuable species like clams and scallops, are under increasing pressure from both anthropogenic and environmental stressors. Over recent decades, shellfish fisheries have suffered dramatic declines due to overexploitation, habitat loss, pollution, and other ecological impacts (Baeta, Breton, Ubach, & Ariza, 2018; Baeta, Ramón, & Galimany, 2014; Cook, 2014, 2019; Edgar & Samson, 2004; Hawes, Lasiak, Smith, & Oengpepa, 2011; Joseph & Jayaprakash, 2003; Morales-Bojórquez, Muciño-Díaz, & Vélez-Barajas, 2008; Peterson, Fegley, & Gaskill, 2008). The striped Venus clam (*Chamelea gallina*), a key fishery resource in the Atlantic-Mediterranean region, illustrates this trend: while Italian landings reached 80,000-100,000 tonnes in the early 1980s, current global catches have fallen below 50,000 tonnes (FAO-GFCM, 2020; Gizzi et al., 2016; Öztürk & Altınok, 2021).

The striped Venus clam (*Chamelea gallina)* is vital to the small-scale fisheries in the Mediterranean and Black Seas, with average annual landings of 47,613 tonnes (FAO-GFCM, 2022). However, many fisheries targeting this species are collapsing, particularly in the western Mediterranean and along the Atlantic coast of Spain (Baeta, Solís, Ballesteros, & Defeo, 2021). In the Gulf of Cadiz, overfishing intensified after the introduction of hydraulic dredgers in 1996, severely depleting clam stocks (Silva et al., 2019). Population resilience is strongly influenced by environmental factors, such as habitat suitability and productivity, with regions like the Alboran Sea supporting higher harvests (García-Fernández et al., 2024). Nevertheless, despite high productivity enabling partial recovery in some areas (Baeta et al., 2021; Llope, 2017), landings have drastically declined: catches along the northwest Mediterranean coast dropped by more than 80% between 1995 to 2004, reaching zero by 2021 (Baeta, unpublished). Ensuring the long-term viability of *C. gallina* fisheries requires effective governance and regulatory framework that align with biological stock boundaries (Baeta et al., 2018). Adaptive management strategies offer a pathway to recovery, emphasizing the need for comprehensive approaches to mitigate overfishing and environmental degradation (Dağtekin & Özyurt, 2023).

Recent population genetics studies of this species have been conducted across various regions of their global distribution, utilising putatively neutral markers such as microsatellites, mtDNA haplotype diversity, and Single Nucleotide Polymorphisms (SNP). Öztürk and Altınok (2021) reported genetic homogeneity within the Black Sea, Levantine Sea, Adriatic Sea, and Atlantic Ocean, which indicate effective panmixia throughout the species’ distribution and suggest that stocks could be managed at a regional scale. In contrast, Carducci et al. (2020) carried out the first population genomics analysis using 2,004 putatively neutral SNPs from samples in the Adriatic and Tyrrhenian Seas. They observed substantial differentiation between the two basins; however, panmixia was observed within each basin. Despite a marked decline of *C. gallina* in the Adriatic Sea over the past two decades, neutral genetic diversity appeared largely preserved, indicating that the population retains significant evolutionary potential. However, adaptive loci were not investigated, while no simple relationship between neutral diversity estimates can be drawn, instead a better understanding of adaptive and functional genetic diversity is necessary to evaluate population health (Teixeira & Huber, 2021).

Here, we explicitly test whether major Mediterranean biogeographic boundaries and environmental gradients jointly shape connectivity and local adaptation in the striped Venus clam (*Chamelea gallina*), and whether these patterns delineate biologically meaningful management units. Specifically, we hypothesise that (i) high dispersal potential leads to widespread connectivity and weak population structure at putatively neutral loci across much of the Mediterranean, while (ii) spatially heterogeneous selection associated with basin-scale environmental gradients counteracts gene flow, generating regionally coherent adaptive divergence. We further predict that (iii) key environmental variables, particularly sea surface temperature, salinity and nutrient availability, act as selective axes structuring adaptive genomic variation. By integrating neutral and adaptive SNP data within a seascape genomics framework, this study aims to clarify how Mediterranean seascape heterogeneity shapes evolutionary and biogeographic patterns in an exploited marine bivalve, and to assess the implications of these patterns for population resilience and spatially explicit management.

## 2 Materials and Methods

### 2.1 Sampling design and filtering steps

A total of 226 *C. gallina* specimens were genotyped using the DArTseq™ platform. To ensure high SNP data quality, we implemented a sequence of filtering procedures based on the quality metrics provided by the DArTseq™ pipeline. SNP filtering was conducted in R using the dartR (Gruber, Unmack, Berry, & Georges, 2018) and poppr (Kamvar, Tabima, & Grünwald, 2014) packages.

Loci were selected according to the following criteria: a minimum allele frequency (MAF > 0.05), a missing data rate below 35% per locus and per individual, and a pairwise linkage disequilibrium threshold of R² < 0.8 (refer to Table S1). We filtered individuals based on heterozygosity, using a threshold of 0.5. To reduce potential biases in population structure inference, pairwise relatedness was estimated: full siblings (r^2^ > 0.4) and half-sibling (r^2^ > 0.2) were excluded using the R package related (Pew, Muir, Wang, & Frasier, 2015). A neutral dataset was then created by removing loci that significantly departed from Hardy-Weinberg equilibrium (HWE) in more than 70% of sampling sites, using the *gl.filter*.hwe function in dartR. This step was performed after outlier detection to avoid excluding loci potentially under selection, which often deviate from HWE. This filtering pipeline produced a high-quality SNP dataset with strong repeatability, suitable for downstream population genomic analyses.

### 2.2 Outlier detection

Outlier loci were detected using two complementary approaches. The first method employed was pcadapt (Luu, Bazin, & Blum, 2017), which identifies loci potentially under selection by analysing individual-level genotyping data. This approach relies on principal component analysis (PCA) to characterise the main axes of genetic variation and identifies SNPs showing significant associations with these axes, indicative of putative adaptive divergence. For each SNP, a q-value was computed to account for multiple testing and control the false discovery rate. SNPs with a q-value below 0.05 were considered outliers and interpreted as candidates for loci under selection.

The second method employed was OutFLANK, an approach implementing the framework developed by Whitlock and Lotterhos (2015). OutFLANK detects loci potentially under selection by estimating the distribution of F_ST_ among populations (here defined as sampling locations) expected under neutrality. This method fits a probability model to a truncated distribution of observed F_ST_ values, deliberately excluding extreme values that are likely influenced by selection. For each locus, q-values were computed based on the inferred neutral distribution. Loci with q-values below 0.05 were considered outliers and interpreted as candidates for spatially heterogeneous selection across sampling locations. This significant threshold was consistent with that used in the pcadapt analysis.

### 2.3 Population structure at both neutral and outlier loci

Population structure was investigated using both neutral and adaptive SNP datasets through two complementary clustering approaches. First, a non-model-based *k*-means clustering method was applied using the *find.cluster* function from the adegenet R package (Jombart, 2008). This approach identifies the optimal number of genetic clusters (*K*) by evaluating the Bayesian Information Criterion (BIC) across increasing values of *K*. Prior to clustering, genetic data were transformed using PCA, and clustering was performed without assuming any explicit population genetic model.

Second, a model-based clustering analysis was conducted using sparse non-negative matrix factorisation (sNMF) as implemented in the LEA R package (Frichot & François, 2015). Similar to STRUCTURE (Pritchard, Stephens, & Donnelly, 2000), this approach estimates individual ancestry coefficients and infers population structure under the assumption of HWE. The optimal number of clusters was determined using the cross-entropy criterion, which evaluates genotype imputation accuracy through cross-validation, with lower values indicating a better model fit. Values of *K* ranging from one to twelve (i.e., *n* + 1, where *n* corresponds to the number of sampling locations) were tested with 10 replicate runs performed for each *K*, using a regularisation parameter of α = 1,000. The *K* value associated with the lowest cross-entropy score was retained as the best estimate of the number of ancestral populations.

### 2.4 Signatures of local adaptation

Environmental variables, including sea surface temperature (SST), sea surface salinity (SSS), chlorophyll concentration, nitrate, phosphate, dissolved oxygen, iron, pH, and phytoplankton biomass, were extracted from the Bio-ORACLE database for the available temporal range (2000–2014) (Assis et al., 2018; Tyberghein et al., 2012). Associations between genetic variations and environmental gradients were investigated using redundancy analysis (RDA) with SNP allele frequencies as response variables and environmental predictors as explanatory variables.

Prior to model fitting, multicollinearity among predictors was assessed and only variables with a variance inflation factor (VIF) below 10 were retained. The significance of RDA axes was evaluated using permutation tests. SNP loadings on significant axes were extracted and standardized into *z*-scores, from which p-values were derived and corrected for multiple testing using the false discovery rate. SNPs located in the upper and lower tails of the corrected *z*-score distributions were considered candidate outliers potentially under selection.

To account for the effects of demographic structure and reduce false positives associations, a partial RDA was additionally performed. In this analysis, individual principal component scores derived from the neutral SNP dataset were included as conditioning variables (Capblancq, Luu, Blum, & Bazin, 2018). This approach allows the detection of genotype-environment associations while controlling for background population structure (Forester, Lasky, Wagner, & Urban, 2018).

## 3 Results

### 3.1 Genotyping

We successfully genotyped 1,293 SNPs in 226 individuals and applied two outlier detection methods without a priori assumption regarding the selective agent: pcadapt and OutFLANK. Pcadapt identified 213 SNPs exhibiting signatures of selection, whereas OutFLANK detected no outliers, resulting in no overlap between the two approaches. Population structure analyses of the full SNP dataset revealed three genetic clusters (Adriatic Sea, Gulf of Cadiz, and all other locations; Figure S1), which were used to determine the number of principal components (*K*) in the pcadapt analysis. The dataset was subsequently partitioned into putatively neutral (1,080 SNPs) and putatively adaptive loci (213 SNPs).

### 3.2 Population structure

Using the neutral genome-wide SNPs, no evidence of genetic clustering was detected with the *k*-means method, with the best-supported number of clusters being *K* = 1 (BIC = 1012.8). When *K* was forced to two, subtle structure emerged (CV = 1014.4; Figure S2), reflecting differentiation between the Adriatic Sea (AS) and the remaining sites. Model-based clustering using LEA yielded consistent results, identifying three clusters corresponding to the Adriatic Sea (AS), the Gulf of Cadiz (CA) and all other locations combined.

In contrast, analyses based on the putatively adaptive SNPs revealed stronger structure. The *k*-means method supported *K* = 5 according to the BIC, uncovering clear differentiation among regions: Alboran Sea (AL), Adriatic Sea (AS), Gulf of Cadiz (CA), the Balearic sampling sites (DE/GA), and the Tyrrhenian Sea (TS), each forming distinct genetic units (Figure 2), with no temporal differentiation detected among sampling years. Notably, eight individuals sampled in the Balearic region were genetically assigned to Alboran Sea, whereas three individuals from the Alboran Sea were assigned to the Balearic region. LEA analyses confirmed this pattern, identifying four genetic clusters in which the Balearic sites grouped with those from the Alboran Sea (Figure S2).

**Figure 1.**
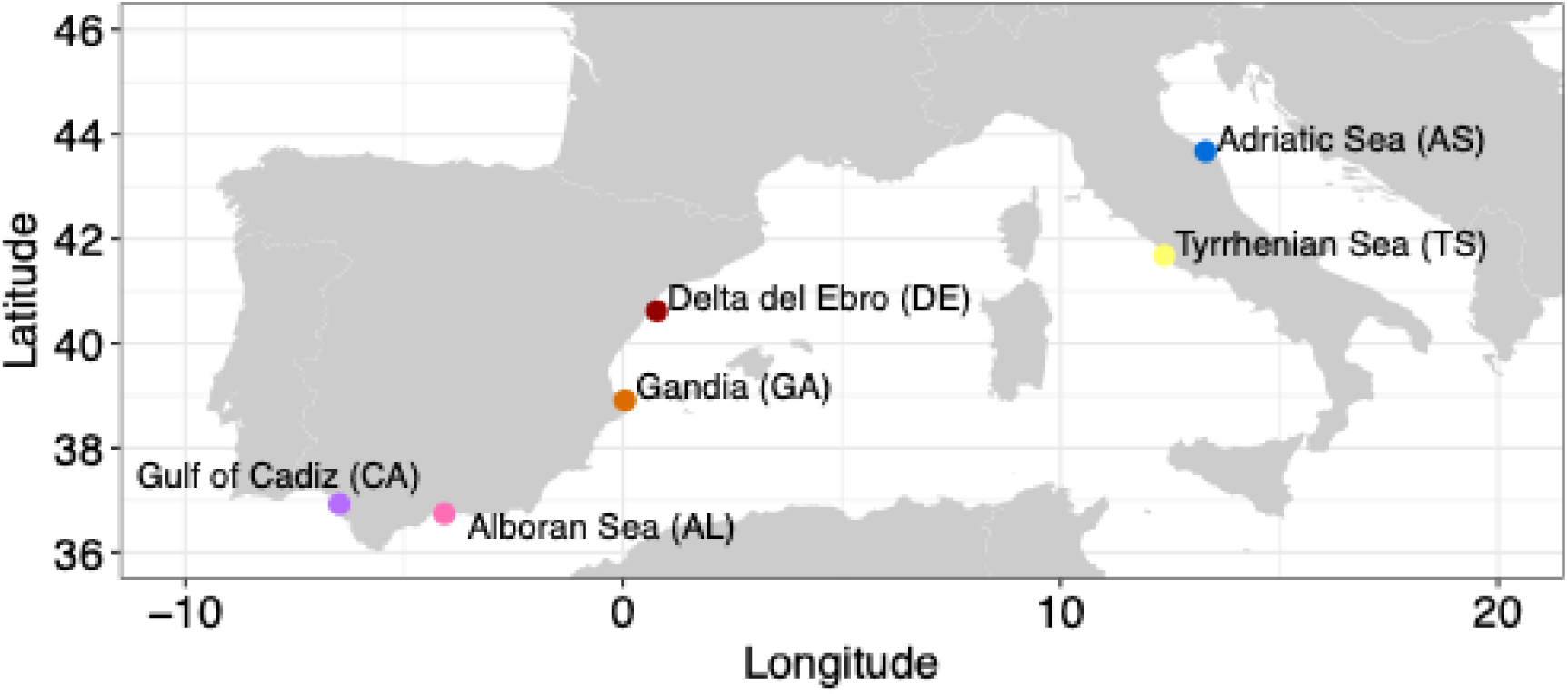
Sampling map.

**Figure 2.**
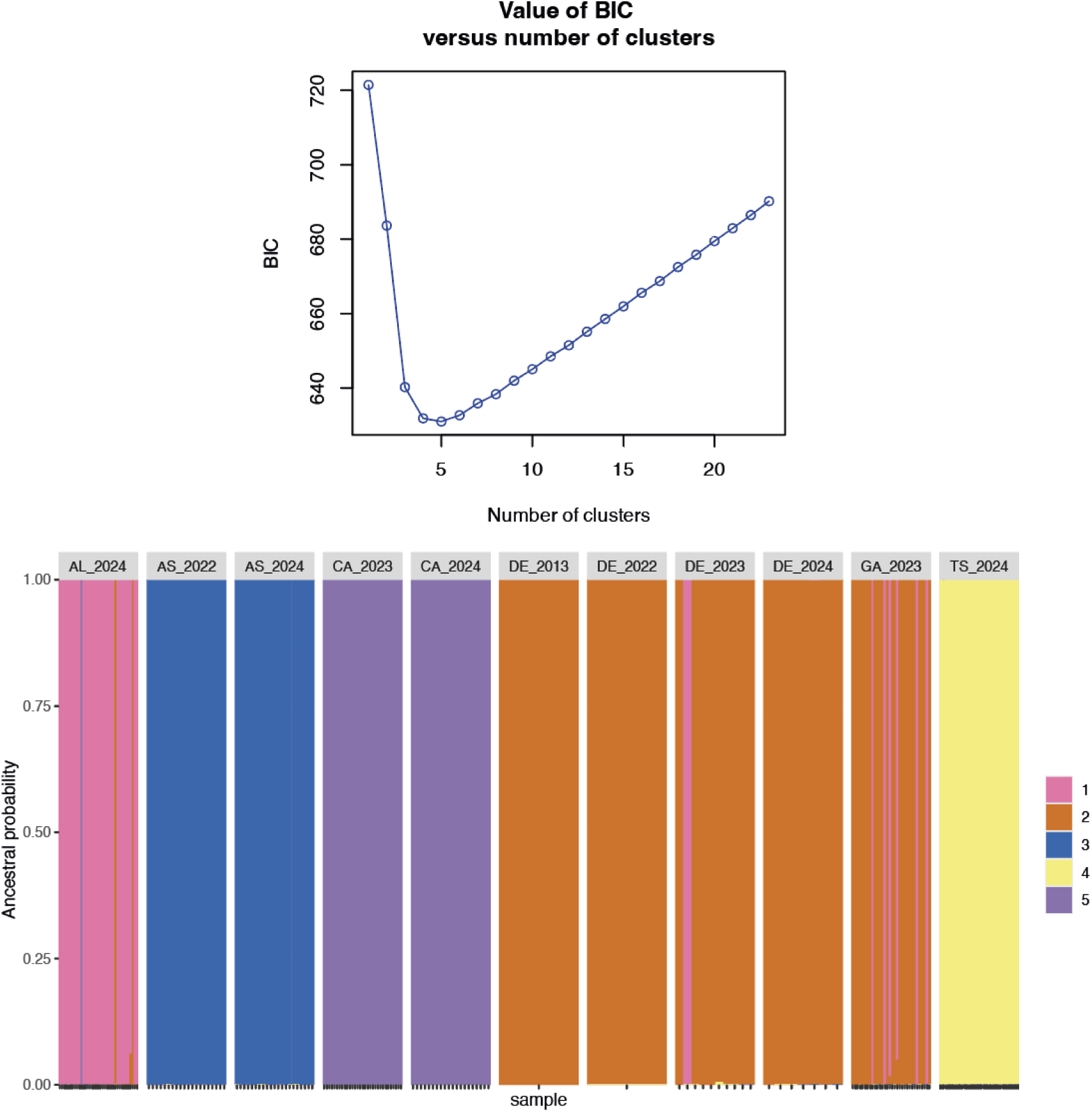

### 3.3 Genetic diversity

Pairwise F_ST_ estimates ranged from 0.0027 to 0.0515 (Figure 3) with the highest differentiation observed between the Adriatic Sea (AS) and the Tyrrhenian Sea (TS), and the lowest between the Alboran Sea (AL) and Gandia (GA). All pairwise comparisons were statistically significant, except for AL-GA and AL-DE. Neutral observed heterozygosity ranged from 0.0517 and 0.1575 and was generally similar across sites, with mean H_O_ of 0.101, 0.104, 0.095 for AL, AS, and DE, respectively, but slightly higher in GA (0.133), CA (0.125) and TS (0.121). Genetic diversity in GA was significantly higher than in all other sites.

**Figure 3.**
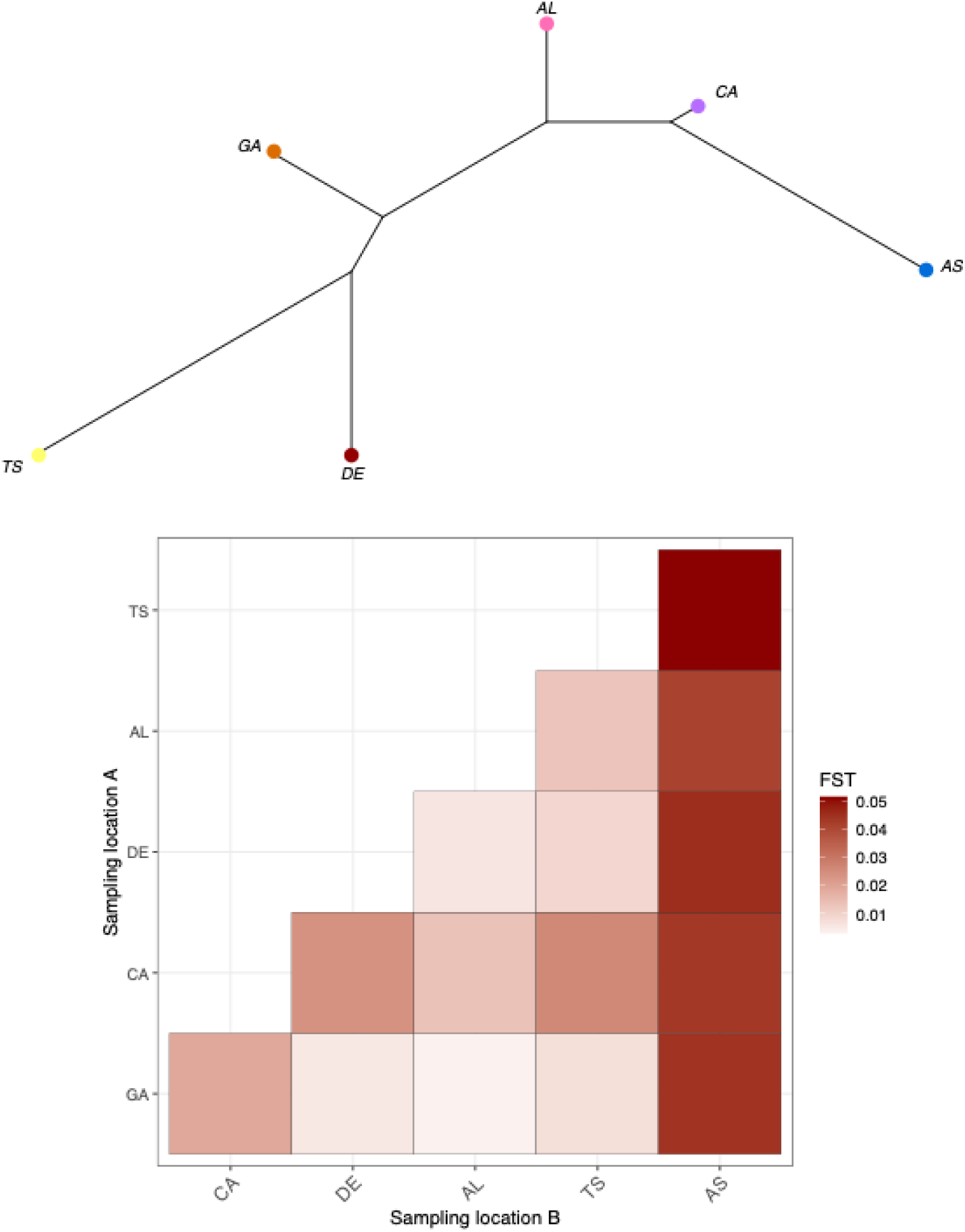
(top) Tree of genetic differentiation unrooted (bottom) heatmap of pairwise FST comparisons across sampling sites. Pairwise FST comparisons are all significant except between GA/DE and AL.

### 3.4 Genotype environment association

Out of the nine environmental variables initially considered, three were retained after assessing pairwise correlation and multicollinearity (|r|> 0.7 and VIF ≥5): mean SST, mean SSS and mean nitrate. Each variable showing a clear gradient across the Mediterranean sampling sites.

Although the redundancy analysis explained a relatively small proportion of the total genetic variance (< 1 %), permutation tests confirmed that the axes were statistically significant. Low variance explained is common in large SNP datasets, where most loci are likely neutral and only a subset respond to environmental gradients. Despite the modest overall variance, the significant RDA axes allowed the identification of SNPs with extreme loadings, which were considered candidate outliers potentially under selection. All three retained environmental variables were significant (p = 0.001).

A total of 19 and 9 SNPs was identified as candidate adaptive markers along the first and second significant axes. The first axis explained 0.9% of the constrained variance, and the second explained 0.5%. The RDA biplot illustrates variation across sampling locations with changes in nitrate, sea surface temperature and salinity in relation to population structure (Figure 4).

**Figure 4.**
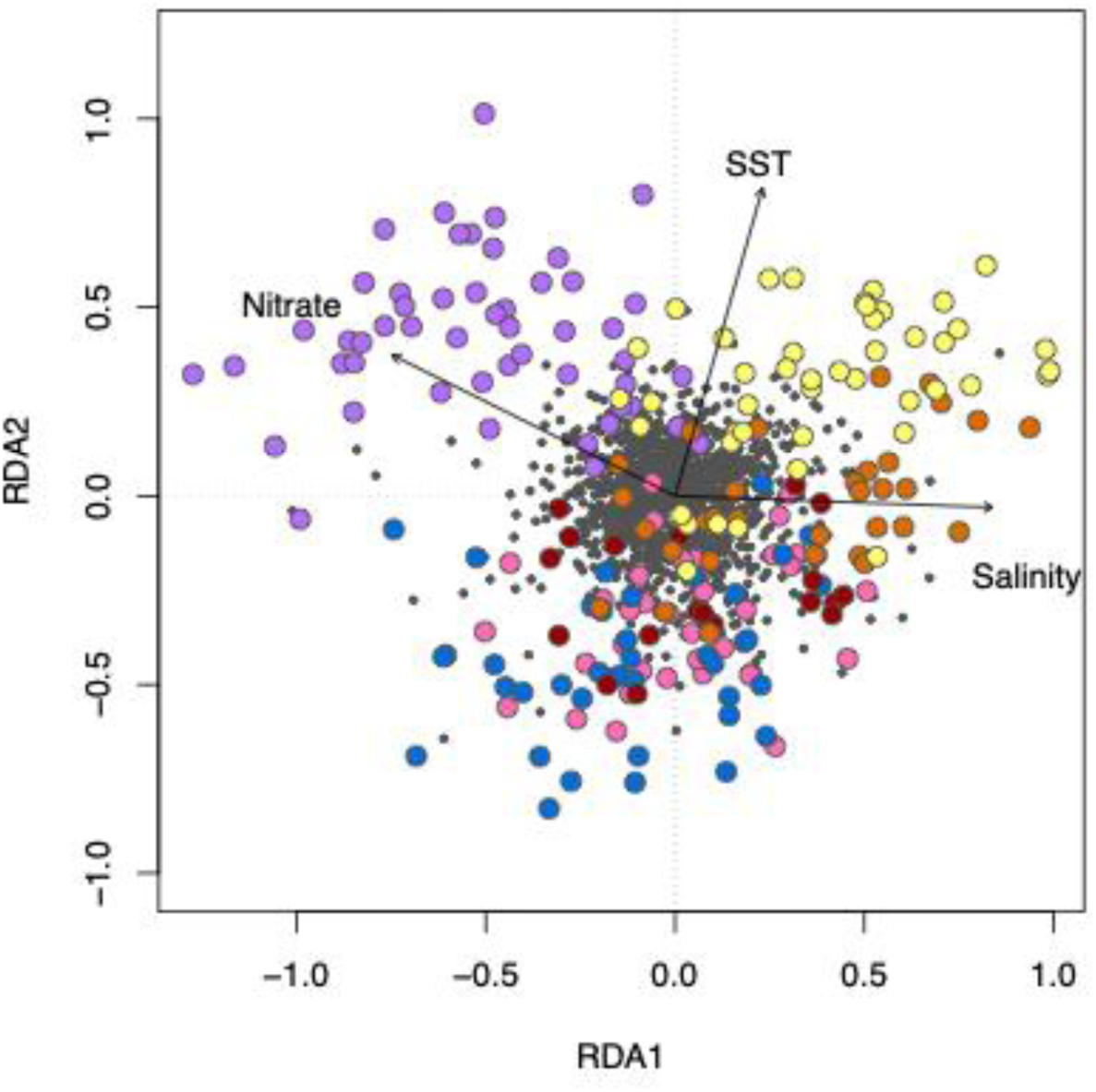
Redundancy Canonical Analysis (RDA) displaying the influence of three environmental variables on individual genomic variation of *Chamelea gallina* from Mediterranean Sea.

## 4 Discussion

### 4.1 Connectivity with gene flow, yet pronounced adaptive structure

This study provides compelling evidence that genetic connectivity and local adaptation coexist in the striped Venus clam (*Chamelea gallina*) across the Mediterranean Sea and adjacent Atlantic waters. Collectively, our analyses address the central expectations outlined in the Introduction by showing widespread connectivity at neutral loci, pronounced regional divergence at adaptive loci, and a key role of basin-scale environmental gradients in shaping this adaptive variation. Using genome-wide SNPs in a seascape framework, we show that populations are largely panmictic at putatively neutral loci, yet display clear and geographically coherent differentiation at potentially adaptive loci. This pattern reflects a growing consensus in marine population genomics: high dispersal potential does not preclude adaptive divergence when strong and persistent environmental gradients impose spatially heterogeneous selection (Benestan, 2020; Gagnaire et al., 2015).

Neutral homogeneity across most Mediterranean sites is consistent with the planktonic larval stage of *C. gallina*, which may last several weeks and facilitates long-distance dispersal via surface currents. Similar discrepancies between neutral and adaptive structure have been documented in a wide range of marine taxa, including sardine (*Sardina pilchardus*), anchovy (*Engraulis encrasicolus*), striped red mullet (*Mullus surmuletus*), and white seabream (*Diplodus sargus*), where adaptive loci reveal strong spatial structuring despite weak neutral differentiation (Alexandridis et al., 2025; Antoniou et al., 2023; Boulanger et al., 2022; Dalongeville, Benestan, Mouillot, Lobreaux, & Manel, 2018). Together, these studies support the view that selection can maintain adaptive divergence even in the presence of gene flow, a process particularly relevant in environmentally complex basins such as the Mediterranean Sea.

### 4.2 Biogeographic breaks and their genomic signatures

The spatial configuration of adaptive genetic clusters in *C. gallina* closely mirrors well-established Mediterranean biogeographic regions, reinforcing the biological relevance of oceanographic fronts and basin-scale discontinuities. The strong differentiation of Adriatic populations, detected at both neutral and adaptive loci, is consistent with the semi-enclosed nature of the Adriatic Sea, its shallow bathymetry, high freshwater input, and distinct circulation patterns, all of which can reduce connectivity with the rest of the Mediterranean (Poulain, 2001; Sefc et al., 2020). Similar Adriatic–Mediterranean breaks have been reported in benthic invertebrates and fishes, often attributed to both historical isolation and contemporary ecological gradients (Marie et al., 2016; T. Patarnello, F. A. M. J. Volckaert, & R. Castilho, 2007).

The Atlantic Gulf of Cádiz population also emerged as a distinct genetic unit, highlighting the long-recognized Atlantic–Mediterranean transition as a partial barrier to dispersal. The Strait of Gibraltar and the Almería–Oran Front (AOF) create sharp contrasts in temperature, salinity, and productivity that limit effective gene flow in many taxa (Galarza et al., 2009; Pascual et al., 2017; Tintore et al., 1988). The differentiation observed here aligns with previous findings in *C. gallina* and other bivalves, such as *Donax trunculus*, where genetic breaks coincide with this transition zone (Marie et al., 2016; Öztürk & Altınok, 2021).

Within the western Mediterranean, adaptive differentiation separating the Alboran, Balearic, Tyrrhenian, and Adriatic regions further emphasizes the role of mesoscale oceanography. Fronts such as the Balearic Front and circulation features associated with the Northern Current can generate retention zones and limit larval exchange, thereby facilitating local adaptation (Balbín et al., 2012; Pinot et al., 1994). The partial assignment of individuals between the Alboran and Balearic regions likely reflects ongoing gene flow modulated by these dynamic processes, resulting in porous rather than absolute barriers, a hallmark of Mediterranean biogeography.

### 4.3 Environmental drivers of adaptive divergence

Genotype–environment association analyses identified sea surface temperature, salinity, and nitrate concentration as key predictors of adaptive genetic variation in *C. gallina*. These variables represent fundamental axes of environmental heterogeneity in the Mediterranean and are known to influence bivalve physiology, growth, and survival. Temperature affects metabolic rates and reproductive timing, salinity shapes osmoregulatory processes, and nutrient availability determines primary productivity and food supply for suspension feeders (García-Fernández et al., 2024; Gizzi et al., 2016).

Although the proportion of explained genetic variance was modest, this is expected in polygenic systems where adaptation is driven by many loci of small effect (Capblancq et al., 2018; Forester et al., 2018). Similar effect sizes have been reported in seascape genomic studies of marine fishes and invertebrates, yet these signals consistently correspond to ecologically meaningful gradients (Antoniou et al., 2023; Dalongeville et al., 2018). The congruence between environmental gradients, adaptive loci, and biogeographic regions in *C. gallina* therefore provides strong evidence that local adaptation is actively shaping population structure.

### 4.4 Genetic diversity and population resilience

Despite severe declines in abundance and fishery collapses across much of the species’ range, neutral genetic diversity remained relatively high and homogeneous among populations, including those from historically overexploited regions. This finding echoes previous results from the Adriatic Sea, where genetic diversity persisted despite long-term demographic decline (Carducci et al., 2020). Such patterns are increasingly recognized in marine species with large effective population sizes, where genetic erosion may lag far behind demographic collapse (Hauser, Adcock, Smith, Bernal Ramírez, & Carvalho, 2002; Shaw et al., 2025).

However, as emphasized by recent theoretical and empirical work, neutral genetic diversity alone is a poor proxy for adaptive potential or extinction risk (Teixeira & Huber, 2021). The detection of region-specific adaptive variation in *C. gallina* highlights that the loss of locally adapted populations could erode functional diversity long before neutral diversity declines become apparent. Preserving adaptive differentiation is therefore essential to maintain the evolutionary resilience of exploited marine species, particularly under accelerating climate change.

### 4.5 Implications for fisheries management and conservation

From a management perspective, our results strongly support the need to align management units with adaptive population structure rather than relying solely on neutral connectivity. Current management frameworks for *C. gallina* often operate at broad regional scales, implicitly assuming demographic homogeneity (Baeta et al., 2018; FAO-GFCM, 2022). Our findings indicate that such approaches risk overlooking biologically meaningful population boundaries shaped by local adaptation.

Adaptive divergence across biogeographic regions suggests that one-size-fits-all management strategies may be ineffective or even detrimental, as locally adapted populations may respond differently to exploitation and environmental change. Incorporating genomic information into fisheries governance, as advocated for other marine resources, could enable more targeted regulations that preserve both demographic viability and adaptive capacity (Benestan, 2020; Fuentes-Pardo, Farrell, Pettersson, Sprehn, & Andersson, 2023).

In the context of climate change, maintaining adaptive diversity across environmental gradients will be critical. The Mediterranean Sea is warming faster than the global ocean, with projected increases in temperature and altered nutrient regimes likely to intensify selection pressures (Bianchi et al., 2012). Protecting regionally adapted populations of *C. gallina* may therefore enhance the species’ capacity to persist under future conditions.

### 4.6 Broader biogeographic implications

Beyond the focal species, our results reinforce a central paradigm in marine biogeography: that contemporary biodiversity patterns emerge from the interplay between historical basin configuration, persistent oceanographic features and spatially structured selection. The Mediterranean Sea, often considered highly connected, harbours fine-scale adaptive mosaics that remain invisible to neutral markers alone. Incorporating adaptive genomic data into biogeographic frameworks is crucial for refining regionalization schemes, forecasting species responses to climate-driven environmental change, and embedding evolutionary processes into conservation planning across semi-enclosed seas.

### 4.7 Conclusion

This study explicitly set out to test whether major Mediterranean biogeographic boundaries and environmental gradients shape genomic connectivity and local adaptation in *Chamelea gallina*, and to assess whether these patterns are relevant for defining biologically meaningful management units. Our results directly fulfil these aims. First, we show that gene flow across much of the Mediterranean is sufficient to homogenise neutral genetic variation, consistent with the species’ planktonic larval dispersal. Second, we demonstrate that adaptive genomic variation is structured along basin-scale biogeographic divisions and environmental gradients, indicating that selection counteracts dispersal and promotes regionally coherent divergence. Third, we identify sea surface temperature, salinity and nutrient availability as key selective axes underpinning this divergence. Collectively, these findings support our central hypothesis that Mediterranean seascape heterogeneity generates adaptive mosaics that remain undetected by neutral markers alone, and that incorporating this adaptive structure is essential for understanding biogeographic patterns, population resilience and the sustainable management of exploited marine species.

**Table 1.** Filtering steps undertaken with subsequent individuals and SNPs remaining per filtering stage.

|  | SNPs | Individuals |
| --- | --- | --- |
| **Neutral SNPs** |  |  |
| Raw dataset | 20,626 | 273 |
| MAF > 0.05 | 7,685 | 273 |
| Loci with less than 40% of missing | 1,293 | 273 |
| Individuals with less than 40% of missing | 1,293 | 226 |
| Hardy-Weinberg Equilibrium in at 70% of the sampling locations (P-value < 0.05) |  |  |
| LD < 0.2 | 1,293 | 226 |
| Removal of related individuals | 1,293 | 226 |
| Removal of outliers | **1,080** | 226 |
| **Outlier SNPs** |  |  |
| PCAdapt | 213 |  |
| OUTFLANK | 0 |  |

**Table 2.**
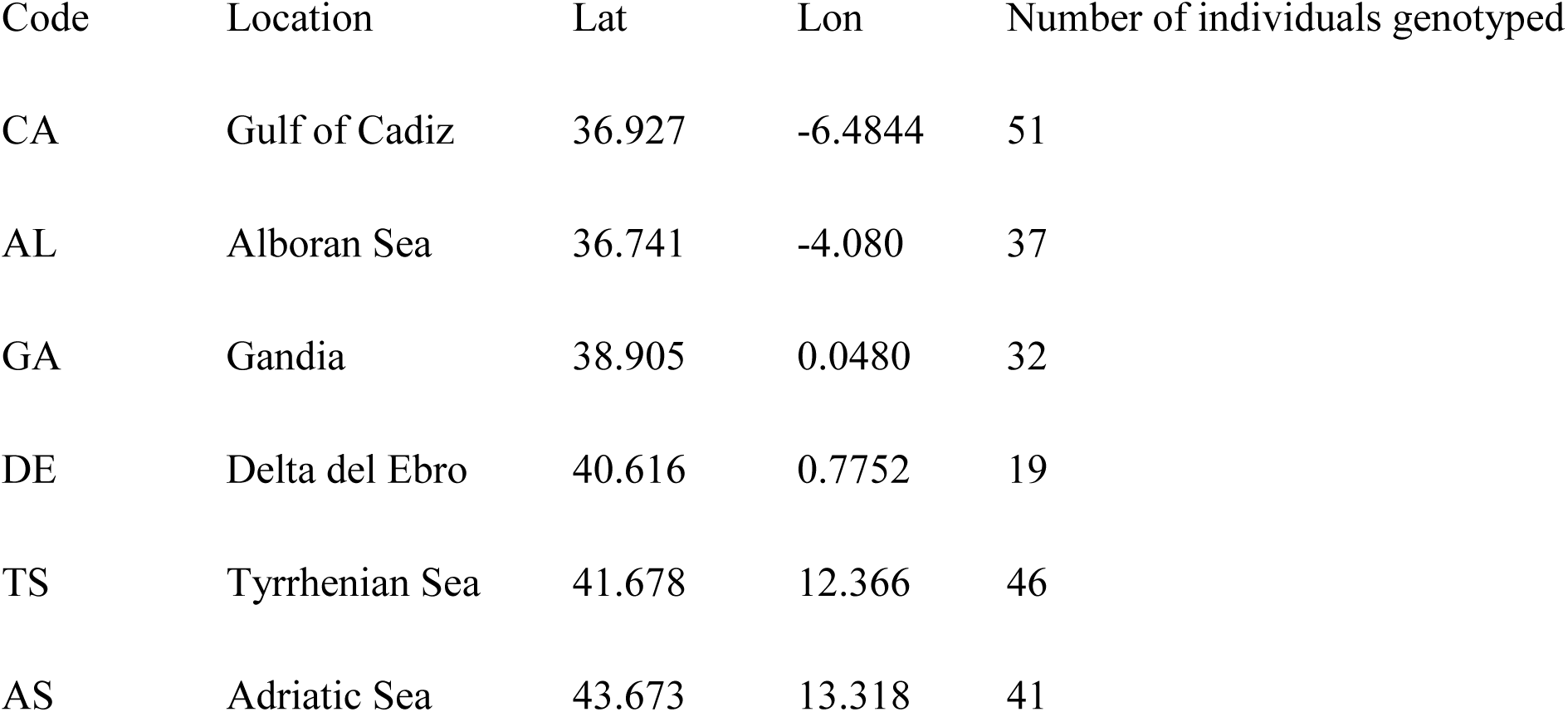
Sampling sites.

| Code | Location | Lat | Lon | Number of individuals genotyped |
| --- | --- | --- | --- | --- |
| CA | Gulf of Cadiz | 36.927 | -6.4844 | 51 |
| AL | Alboran Sea | 36.741 | -4.080 | 37 |
| GA | Gandia | 38.905 | 0.0480 | 32 |
| DE | Delta del Ebro | 40.616 | 0.7752 | 19 |
| TS | Tyrrhenian Sea | 41.678 | 12.366 | 46 |
| AS | Adriatic Sea | 43.673 | 13.318 | 41 |

## 5 Acknowledgments

We are grateful to Department d′Agricultura, Ramaderia i Pesca de la Generalitat de Catalunya, Conselleria d’Agricultura, Aigua, Ramaderia i Pesca de la Generalitat Valenciana, Consejería de Agua, Agricultura, Ganadería y Pesca de la Región de Murcia and Consejería de Agricultura, Pesca, Agua y Desarrollo Rural (Junta de Andalucía) for facilitating sample collection.

---

**Figure S1.**
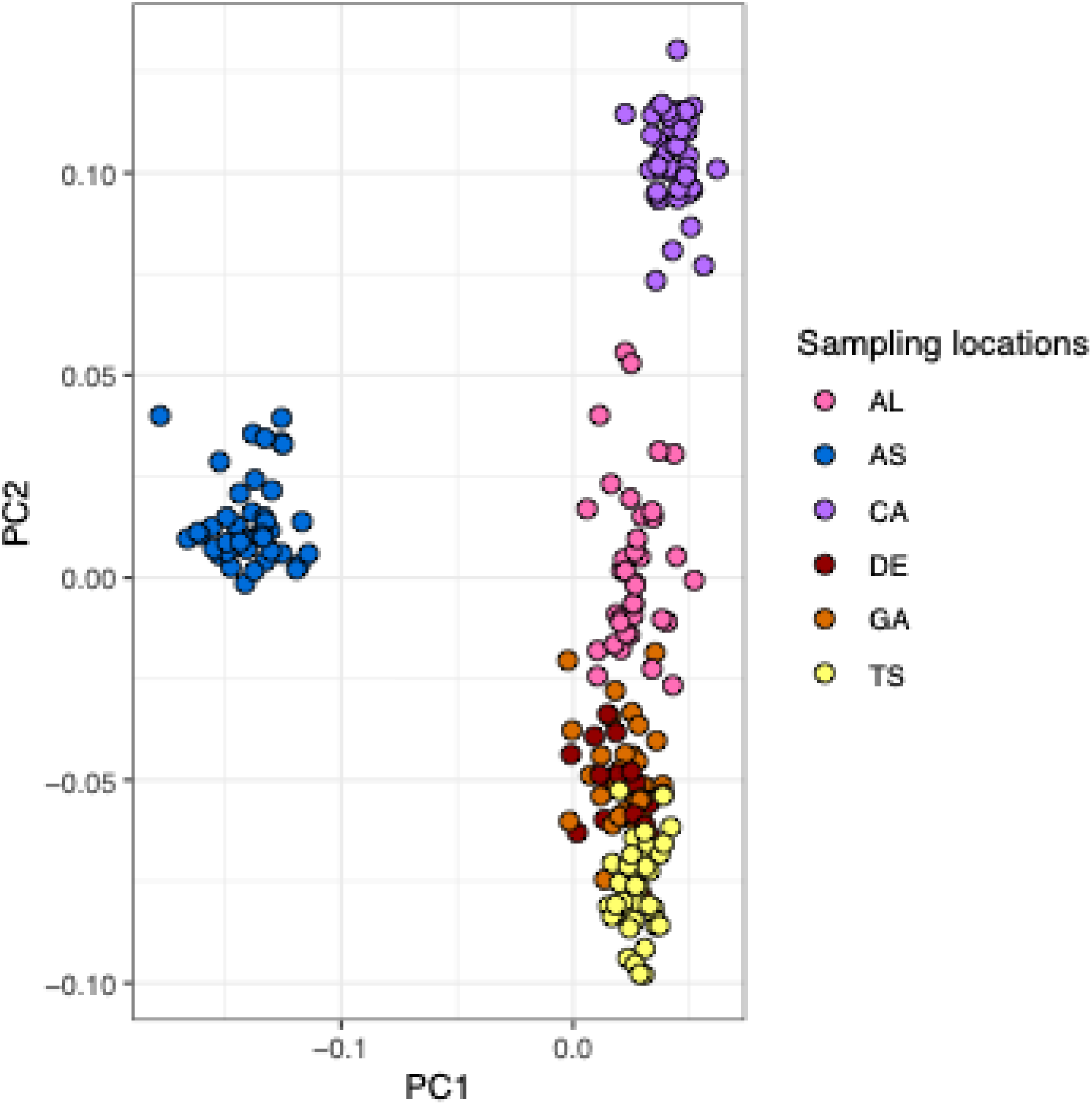
PCAdapt figure showing K = 3.

**Figure S2.**
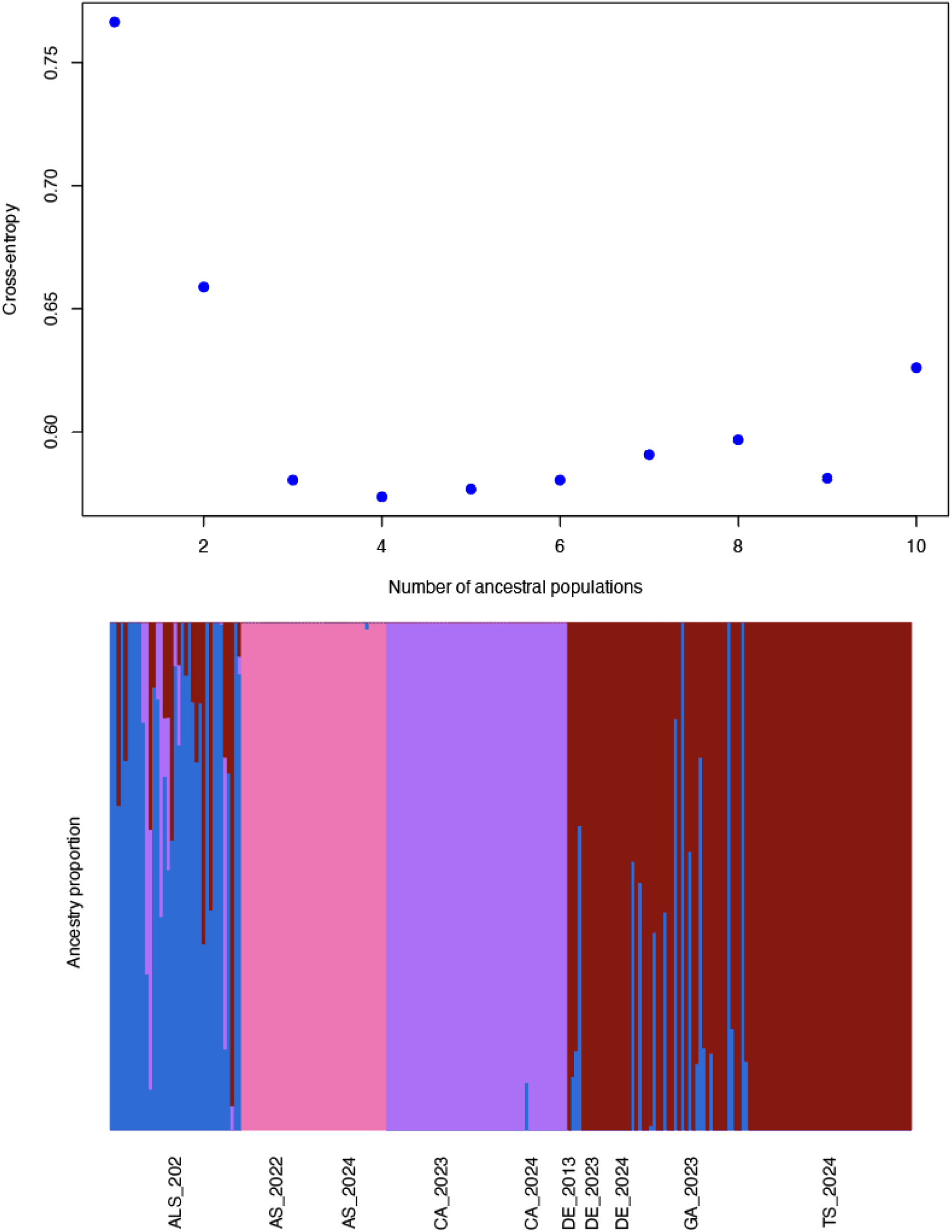

